# Does women’s interpersonal anxiety track changes in steroid hormone levels?

**DOI:** 10.1101/456319

**Authors:** Amanda C Hahn, Lisa M DeBruine, Lola A Pesce, Andrew Diaz, Christopher L Aberson, Benedict C Jones

## Abstract

Findings for progesterone and anxiety in non-human animals led to the hypothesis that women’s interpersonal anxiety will track changes in progesterone during the menstrual cycle. There have been few direct tests of this hypothesis, however. Consequently, we used a longitudinal design to investigate whether interpersonal anxiety (assessed using the anxious jealousy subscale of the relationship jealousy questionnaire) tracked changes in salivary steroid hormones during the menstrual cycle in a large sample of young adult women (N=383). We found no evidence for within-subject effects of progesterone, estradiol, their interaction or ratio, testosterone, or cortisol on anxious jealousy. There was some evidence that other components of jealousy (e.g., reactive jealousy) tracked changes in women’s cortisol, however. Collectively, these results provide no evidence for the hypothesis that interpersonal anxiety tracks changes in progesterone during the menstrual cycle.

## Introduction

Inspired by work linking progesterone to stress and anxiety in non-human animals (e.g., rodents, Barbaccia et al., 1996), several studies have investigated whether interpersonal stress and anxiety are related to progesterone in women. For example, Wirth and Schultheiss (2006) reported positive effects of an interpersonal stressor (fear of social rejection) on progesterone in a sample of 37 women and 48 men. Similarly, Gonda et al. (2008) found that women (N=63) reported greater interpersonal sensitivity and anxiety in the luteal phase of the menstrual cycle (when progesterone is typically high) compared to the follicular phase (when progesterone is typically low). More recent work did not replicate this effect of cycle phase on anxiety, however (Reynolds et al., 2018 Study 1).

A potentially important limitation of the studies described above is that they do not show a link between naturally occurring variation in progesterone and interpersonal stress or anxiety (see Reynolds et al., 2018). For example, Gonda et al. (2008) compared interpersonal anxiety in different menstrual cycle phases, but did not directly relate changes in anxiety to changes in measured progesterone. To address this issue, Reynolds et al. (2018 Study 2) found that women’s (N=61) responses on the attachment anxiety subscale of the revised Experiences in Close Relationships questionnaire (Fraley et al., 2000) tracked changes in women’s salivary progesterone during the menstrual cycle. Reynolds et al. (2018 Study 2) reported that attachment anxiety (a form of interpersonal anxiety regarding romantic relationships) increased when progesterone was high. Reynolds et al. (2018) suggested that this progesterone-linked increase in interpersonal anxiety could function to maintain the strength of romantic relationships when raised progesterone prepares the body for pregnancy.

The current study reports a conceptual replication of Reynolds et al. (2018 Study 2). Specifically, we tested whether women’s (N=369) responses on Buunk’s (1997) relationship jealousy questionnaire tracked changes in their salivary steroid hormone levels. Buunk’s (1997) relationship jealousy questionnaire has three subscales: anxious jealousy, possessive jealousy, and reactive jealousy. Anxious jealousy refers to cognitively generated experiences of anxiety, worry, and distrust, which relate to one’s partner’s infidelity. Possessive jealousy refers to the degree of effort an individual invests to prevent their partner from coming into contact with opposite-sex individuals. Reactive jealousy refers to the degree to which an individual experiences negative emotions as a result of their partner’s emotional or sexual infidelity. Given Reynolds et al’s (2018 Study 2) results for attachment anxiety and progesterone, we predicted that women would report greater anxious jealousy when progesterone was high.

### Participants

We tested 383 women (mean age=21.69 years, SD=3.13 years) who reported that they were not using any form of hormonal contraceptive (i.e., reported having natural menstrual cycles). Participants completed up to three blocks of test sessions. Each of the three blocks of test sessions consisted of five weekly test sessions. Women participated as part of a large study of possible effects of steroid hormones on women’s behavior (Jones et al., 2018a). The data analyzed here are all responses from blocks of test sessions where women were not using any form of hormonal contraceptive and both provided saliva samples and completed Buunk’s (1997) relationship jealousy questionnaire. Following these restrictions, 250 women had completed five or more test sessions and 370 of these women completed ten test sessions. One hundred and twenty women completed fewer than five test sessions.

### Procedure

In each test session, women reported their current romantic partnership status (partnered or unpartnered), provided a saliva sample, and completed Buunk’s (1997) relationship jealousy questionnaire.

Buunk’s (1997) relationship jealousy questionnaire consists of three 5-item subscales, each assessing a different component of relationship jealousy. These components are anxious jealousy (M=10.77, SD=5.13), possessive jealousy (M=9.83, SD=4.23), and reactive jealousy (M=18.55, SD=4.02). An example of an anxious jealousy item is “I worry that my partner might leave me for someone else”. An example of a possessive jealousy item is “I demand from my partner that he does not look at other women”. An example of a reactive jealousy item is “How would you feel if your partner discussed personal things with someone else?” Participants responded to each item using 5-point scales, on which higher scores indicate greater relationship jealousy. Following previous research using this questionnaire (e.g., Cobey et al., 2012), participants who were not currently in a romantic relationship at the time of testing were instructed to think back to their last relationship when responding.

### Saliva samples

Participants provided a saliva sample via passive drool (Papacosta & Nassis, 2011) in each test session. Participants were instructed to avoid consuming alcohol and coffee in the 12 hours prior to participation and avoid eating, smoking, drinking, chewing gum, or brushing their teeth in the 60 minutes prior to participation. Each woman’s test sessions took place at approximately the same time of day to minimize effects of diurnal changes in hormone levels (Veldhuis et al., 1988; Bao et al., 2003).

Saliva samples were frozen immediately and stored at −32°C until being shipped, on dry ice, to the Salimetrics Lab (Suffolk, UK) for analysis, where they were assayed using the Salivary 17β-Estradiol Enzyme Immunoassay Kit 1-3702 (M=3.41 pg/mL, SD=1.31 pg/mL, sensitivity=0.1 pg/mL, intra-assay CV=7.13%, inter-assay CV=7.45%), Salivary Progesterone Enzyme Immunoassay Kit 1-1502 (M=149.09 pg/mL, SD=94.12 pg/mL, sensitivity=5 pg/mL, intra-assay CV=6.20%, inter-assay CV=7.55%), Salivary Testosterone Enzyme Immunoassay Kit 1-2402 (M=87.12 pg/mL, SD=26.83 pg/mL, sensitivity<1.0 pg/mL, intra-assay CV=4.60%, inter-assay CV=9.83%), and Salivary Cortisol Enzyme Immunoassay Kit 1-3002 (M=0.23 μg/dL, SD=0.17 μg/dL, sensitivity<0.003 μg/dL, intra-assay CV=3.50%, inter-assay CV=5.08%).

Hormone levels more than three standard deviations from the sample mean for that hormone or where Salimetrics indicated levels were outside the sensitivity range of their relevant ELISA were excluded from the dataset (~1% of hormone measures were excluded for these reasons). The descriptive statistics given above do not include these excluded values. Values for each hormone were centered on their subject-specific means to isolate effects of within-subject changes in hormones. They were then scaled so the majority of the distribution for each hormone varied from -.5 to .5 to facilitate calculations in the linear mixed models. Since hormone levels were centered on their subject-specific means, women with only one value for a hormone could not be included in these analyses.

### Analyses

Linear mixed models were used to test for possible effects of hormonal status on jealousy. Analyses were conducted using R version 3.3.2 (R Core Team, 2016), with lme4 version 1.1-13 (Bates et al., 2014) and lmerTest version 2.0-33 (Kuznetsova et al., 2013). The dependent variable was subscale score (separate models were run for each jealousy subscale). Predictors were scaled and centered hormone levels. Random slopes were specified maximally following Barr et al. (2013) and Barr (2013). Full model specifications and full results for each analysis are given in our Supplemental Information. Data files and analysis code are publicly available at https://osf.io/cjyv9/.

## Results

Scores for each jealousy subscale were analyzed separately. For each subscale we ran three models. The first model (Model 1) included estradiol, progesterone, and their interaction as predictors. The second model (Model 2) included estradiol, progesterone, and estradiol-to-progesterone ratio as predictors. We tested for combined effects of estradiol and progesterone by including the estradiol by progesterone interaction (Model 1) and estradiol-to-progesterone ratio (Model 2) because both approaches have recently been used to test for combined effects of estradiol and progesterone in the hormones and behavior literature (see Puts et al., 2013 and Roney & Simmons, 2013 for examples of studies using one of these two approaches). The third model (Model 3) included only testosterone and cortisol as predictors. Our analysis strategy is identical to that used in Jones et al. (2018a), Jones et al. (2018b), and Jones et al. (2018c) to investigate the hormonal correlates of within-woman changes in mate preferences, disgust sensitivity, and sexual desire, respectively.

### Anxious jealousy

Model 1 showed no significant effects (all absolute beta<2.98, all absolute t<1.61, all p>0.10). Neither Model 2 (all absolute beta<0.25, all absolute t<1.31, all p>0.18) nor Model 3 (both absolute beta<0.20, both absolute t<0.74, both p>0.46) showed any significant effects.

An additional Bayesian analysis was conducted for Models 1 and 2, given the theoretical predictions regarding progesterone. The calculated Bayes factors express a ratio of the likelihood H0 relative to H1 given the data (i.e., values larger than 1 are in favour of H0, assuming that H0 and H1 are equally likely). For Model 1 (which includes estradiol, progesterone, and their interaction as predictors of anxious jealousy), the BF01 was 18166, suggesting that these data are 18166 times more likely to be observed under the null hypothesis. For Model 2 (which includes estradiol, progesterone, and their ratio as predictors of anxious jealousy), the BF01 was 27312, suggesting that these data are 27312 times more likely to be observed under the null hypothesis.

### Possessive jealousy

Model 1 showed no significant effects (all absolute beta<0.07, all absolute t<0.21, all p>0.17). Neither Model 2 (all absolute beta<0.16, all absolute t<0.49, all p>0.19) nor Model 3 (both absolute beta<0.03, both absolute t<0.09, both p>0.51) showed any significant effects.

### Reactive jealousy

Model 1 showed a significant interaction between estradiol and progesterone (beta=-4.31, t=-2.61, p= 0.012), but no other effects (both absolute beta<0.01, both absolute t<0.01, both p>0.49). Model 2 showed no significant effects (all absolute beta<-0.09, all absolute t<-0.28, all p>0.51). Model 3 showed a significant positive effect of cortisol (beta=0.73, t=2.10, p= 0.037) and no significant effect of testosterone (beta=-0.18, t=-0.48, p= 0.628).

### Partnership status

Repeating the analyses described above, this time including partnership status in each model (effect coded: −0.5 = unpartnered, +0.5 = partnered) generally did not alter the patterns of significance described above. Full results for each of these analyses are given in our Supplemental Materials. The one exception was Model 1 for anxious jealousy, which showed a marginally significant interaction between estradiol and partnership status (beta=-1.57, t=-2.00, p= 0.046). Since this interaction was not predicted a priori and was only marginally significant, we suggest it is likely to be a false positive. Thus, we did not explore it further.

### Total jealousy

Repeating all of our previous analyses for a total jealousy score (created by summing the three jealousy subscales) showed no significant hormone effects in any models. The one exception was a positive effect of cortisol in the version of Model 3 that included partnership status as a predictor (beta=1.76, t=2.80, p= 0.007). One hundred and forty-one of these women’s data for total jealousy have previously been reported in Hahn et al. (2016). These women’s scores on the individual jealousy subscales were not analyzed for Hahn et al. (2016).

## Discussion

Our longitudinal analyses of the putative relationship between relationship jealousy and steroid hormones showed no evidence that anxious, possessive, or reactive jealousy tracked changes in women’s progesterone, estradiol, their interaction or ratio, or testosterone. We saw some evidence that both reactive and total jealousy tracked changes in cortisol; reactive and total jealousy tended to be greater when cortisol was high. Collectively, these null results, particularly those for anxious jealousy, provide no support for Reynolds et al’s (2018) hypothesis that sex hormones, and progesterone in particular, regulate interpersonal anxiety in women.

Our null results for anxious jealousy and progesterone are at odds with Reynolds et al’s (2018 Study 2) results. Reynolds et al. (2018 Study 2) reported that responses on the attachment anxiety subscale of the revised Experiences in Close Relationships questionnaire (Fraley et al., 2000) tracked changes in women’s salivary progesterone during the menstrual cycle. Our analyses of responses on the anxious jealousy subscale of Buunk’s (1997) relationship jealousy questionnaire showed no evidence for hormonal regulation of interpersonal anxiety. We suggest that this difference in results is unlikely to be due to differences in the scales used to assess interpersonal anxiety, since both questionnaires assess anxiety about romantic relationships and there is substantial conceptual overlap in their items^1^. The difference in our and Reynolds et al’s (2018 Study 2) results is also not due to our study being underpowered compared with Reynolds et al. (2018 Study 2). We tested >6 times the number of women as Reynolds et al. (2018 Study 2). While it is possible that larger changes in progesterone than are typical during the menstrual cycle (e.g., those that occur during pregnancy) are linked to changes in interpersonal anxiety, our findings suggest that the progesterone-linked cyclic changes in interpersonal anxiety reported by Reynolds et al. (2018 Study 2) might not be robust.

In summary, we found no evidence that interpersonal anxiety tracked changes in women’s steroid hormone levels. In particular, we did not replicate Reynolds et al’s (2018 Study 2) finding that interpersonal anxiety tracked changes in progesterone during the menstrual cycle. These findings are in line with a recent study that failed to find any relationship between progesterone levels and either state- or trait-anxiety in women (Graham & Shin, 2018). Thus, our data do not support the proposal that progesterone-linked increases in interpersonal anxiety function to maintain the strength of romantic relationships when raised progesterone prepares women’s bodies for pregnancy.

## Footnotes

1 Consistent with this suggestion, we found that scores on these two measures were strongly correlated in a sample of 76 women, none of whom had participated in the main study (r = .68, p < .001). This correlation is similar to the correlation between anxious jealousy scores in the first and second test sessions in our main study (r = .80, p<.001).

## Notes

This research was supported by a European Research Council grant awarded to BCJ (OCMATE).

#### Summary of Updates

Bayesian analyses have been added

https://osf.io/cjyv9/

## References

Bao, A.M., Liu, R.Y., van Someren, E.J.W., Hofman, M.A., Cao, Y., Zhou, J.N., 2003. Diurnal rhythm of free estradiol during the menstrual cycle. Eur. J. Endocrinol. 148, 227–232.

Barbaccia, M. L., Roscetti, G., Bolacchi, F., Concas, A., Mostallino, M. C., Purdy, R. H., & Biggio, G. (1996). Stress-induced increase in brain neuroactive steroids: Antagonism by abecarnil. Pharmacology Biochemistry and Behavior, 54, 205–210.

Barr, D. J., Levy, R., Scheepers, C., & Tily, H. J. (2013). Random effects structure for confirmatory hypothesis testing: Keep it maximal. Journal of memory and language, 68(3), 255–278.

Barr, D. J. (2013). Random effects structure for testing interactions in linear mixed-effects models. Frontiers in psychology, 4, 328.

Bates, D., Maechler, M., Bolker, B., Walker, S., 2014. lme4: Linear mixed-effects models using Eigen and S4. R package version 1.1–6. http://CRAN.R-project.org/package=lme4

Buunk, B.P., 1997. Personality, birth order and attachment styles as related to various types of jealousy. Pers. Indiv. Differ. 23, 997–1006.

Cobey, K.D., Buunk, A.P., Roberts, S.C., Klipping, C., Appels, N., Zimmerman, Y., et al., 2012. Reported jealousy differs as a function of menstrual cycle stage and contraceptive pill use: A within-subjects investigation. Evol. Hum. Behav. 33, 395–401.

Fraley, R. C., Waller, N. G., & Brennan, K. A. (2000). An item response theory analysis of self-report measures of adult attachment. Journal of Personality and Social Psychology, 78, 350–364.

Gonda, X., Telek, T., Juhasz, G., Lazary, J., Vargha, A., & Bagdy, G. (2008). Patterns of mood changes throughout the reproductive cycle in healthy women without premenstrual dysphoric disorders. Progress in Neuro-Psychopharmacology and Biological Psychiatry, 32, 1782–1788.

Graham, B. M., & Shin, G. (2018). Estradiol moderates the relationship between state-trait anxiety and attentional bias to threat in women. Psychoneuroendocrinology, 93, 82–89.

Hahn, A. C., Fisher, C., Cobey, K. D., DeBruine, L. M. & Jones, B. C. (2016). A longitudinal analysis of women’s salivary testosterone and intrasexual competitiveness. Psychoneuroendocrinology, 64: 117–122.

Jones, B. C., Hahn, A. C., Fisher, C., Wang, H., Kandrik, M., Han, C., Fasolt, V., Morrison, D. K., Lee, A., Holzleitner, I. J., O’Shea, K. J., Roberts, S. C., Little, A. C. & DeBruine, L. M. (2018a). No compelling evidence that preferences for facial masculinity track changes in women’s hormonal status. Psychological Science, 29, 996–1005.

Jones, B. C., Hahn, A. C., Fisher, C., Wang, H., Kandrik, M., Lee, A., Tybur, J. M. & DeBruine, L. M. (2018b). Hormonal correlates of pathogen disgust: Testing the Compensatory Prophylaxis Hypothesis. Evolution and Human Behavior, 39, 166–169

Jones, B. C., Hahn, A. C., Fisher, C., Wang, H., Kandrik, M. & DeBruine, L. M. (2018c). General sexual desire, but not desire for uncommitted sexual relationships, tracks changes in women’s hormonal status. Psychoneuroendocrinology, 88, 153–157.

Kuznetsova, A., Brockhoff, P.B., Christensen, R.H.B., 2013. lmerTest: Tests for random and fixed effects for linear mixed effect models (lmerobjects of lme4 package). R package version 2.0-6. http://CRAN.R-project.org/package=lmerTest

Papacosta, E., Nassis, G.P., 2011. Saliva as a tool for monitoring steroid, peptide and immune markers in sport and exercise science. J. Sci. Med. Sport. 14, 424–434.

Puts, D.A., Bailey, D.H., Cardenas, R.A., Burriss, R.P., Welling, L.L.M., Wheatley, J.R., Dawod, K. 2013. Women’s attractiveness changes with estradiol and progesterone across the ovulatory cycle. Horm. Behav. 63, 13–19.

R Core Team, 2013. R: A language and environment for statistical computing. R Foundation for Statistical Computing, Vienna, Austria. Version 3.1.1, URL http://www.R-project.org/.

Reynolds, T. A., Makhanova, A., Marcinkowska, U. M., Jasienska, G., McNulty, J. K., Eckel, L. A.,… & Maner, J. K. (2018). Progesterone and women’s anxiety across the menstrual cycle. Hormones and Behavior, 102, 34–40.

Roney, J. R., & Simmons, Z. L. (2013). Hormonal predictors of sexual motivation in natural menstrual cycles. Hormones and Behavior, 63, 636–645.

Veldhuis, J.D., Christiansen, E., Evans, W.S., Kolp, L.A., Rogol, A.D., Johnson, M.L., 1988. Physiological profiles of episodic progesterone release during the midluteal phase of the human menstrual cycle: Analysis of circadian and ultradian rhythms, discrete pulse properties, and correlations with simultaneous luteinizing hormone release. J. Clin. Endocrinol. Metab. 66, 414–421.

Wirth, M. M., & Schultheiss, O. C. (2006). Effects of affiliation arousal (hope of closeness) and affiliation stress (fear of rejection) on progesterone and cortisol. Hormones and Behavior, 50, 786–795.

